# Differences in cortical surface area in developmental language disorder

**DOI:** 10.1101/2023.07.13.548894

**Authors:** Nilgoun Bahar, Gabriel J Cler, Saloni Krishnan, Salomi S Asaridou, Harriet J Smith, Hanna E Willis, Máiréad P Healy, Kate E Watkins

## Abstract

Approximately seven per cent of children have developmental language disorder (DLD), a neurodevelopmental condition associated with persistent language learning difficulties without a known cause. Our understanding of the neurobiological basis of DLD is limited. Here, we used FreeSurfer to investigate cortical surface area and thickness in 54 children and adolescents with DLD and 74 age-matched controls aged 10-16 years. We also examined cortical asymmetries in DLD using an automated surface-based technique. Those with DLD showed smaller surface area bilaterally in the inferior frontal gyrus extending to the anterior insula, in the posterior temporal and ventral occipito-temporal cortex, and in portions of the anterior cingulate and superior frontal cortex. There were no differences in cortical thickness, nor in asymmetry of these cortical metrics. Post-hoc exploratory analyses revealed that surface area in the left fusiform and inferior frontal cortex related to children’s reading and non-word repetition scores, respectively. This study highlights the importance of distinguishing between surface area and cortical thickness in investigating the brain basis of neurodevelopmental disorders and suggests the development of cortical surface area to be of importance to DLD. Future longitudinal studies are required to understand the developmental trajectory of these cortical differences in DLD and how they relate to language maturation.

## 1 INTRODUCTION

Developmental language disorder (DLD) is a neurodevelopmental condition that affects approximately 7.5% of children (Norbury et al., 2016). Children with DLD have persistent difficulties in developing age-appropriate receptive and expressive language skills (Bishop, 2017) and are at increased risk of behavioural, social, academic, and later vocational problems (e.g. Conti-Ramsden et al., 2018; Goh et al., 2021). Importantly, these language deficits cannot be attributed to known biomedical or environmental causes, such as brain injury, neurological conditions (e.g. epilepsy), childhood hearing loss, or social disadvantage (Bishop et al., 2017). Our understanding of the neurobiological basis of DLD remains limited. Here, we used analysis of brain images obtained in a large, well-characterised cohort of children with DLD and typically developing (TD) controls to explore for differences in cortical morphometry that could be related to differences in language learning.

The existing literature on the neuroanatomical correlates of DLD have used manual tracing or semi-automated methods to study the size, shape, or asymmetry of cortical and subcortical structures. Cumulative evidence indicates a distributed, bilateral pattern of subtle morphological abnormalities focused on perisylvian cortical areas involved in speech and language, and subcortical structures implicated in habit formation and sequential learning (see Mayes et al., 2015 for a review). However, findings are inconsistent among these studies in terms of the location and direction of differences (i.e. whether regions are larger or smaller in DLD) possibly due to different methods used, small sample sizes, and the age ranges of different cohorts (see Table 1 for details).

**Table 1.**
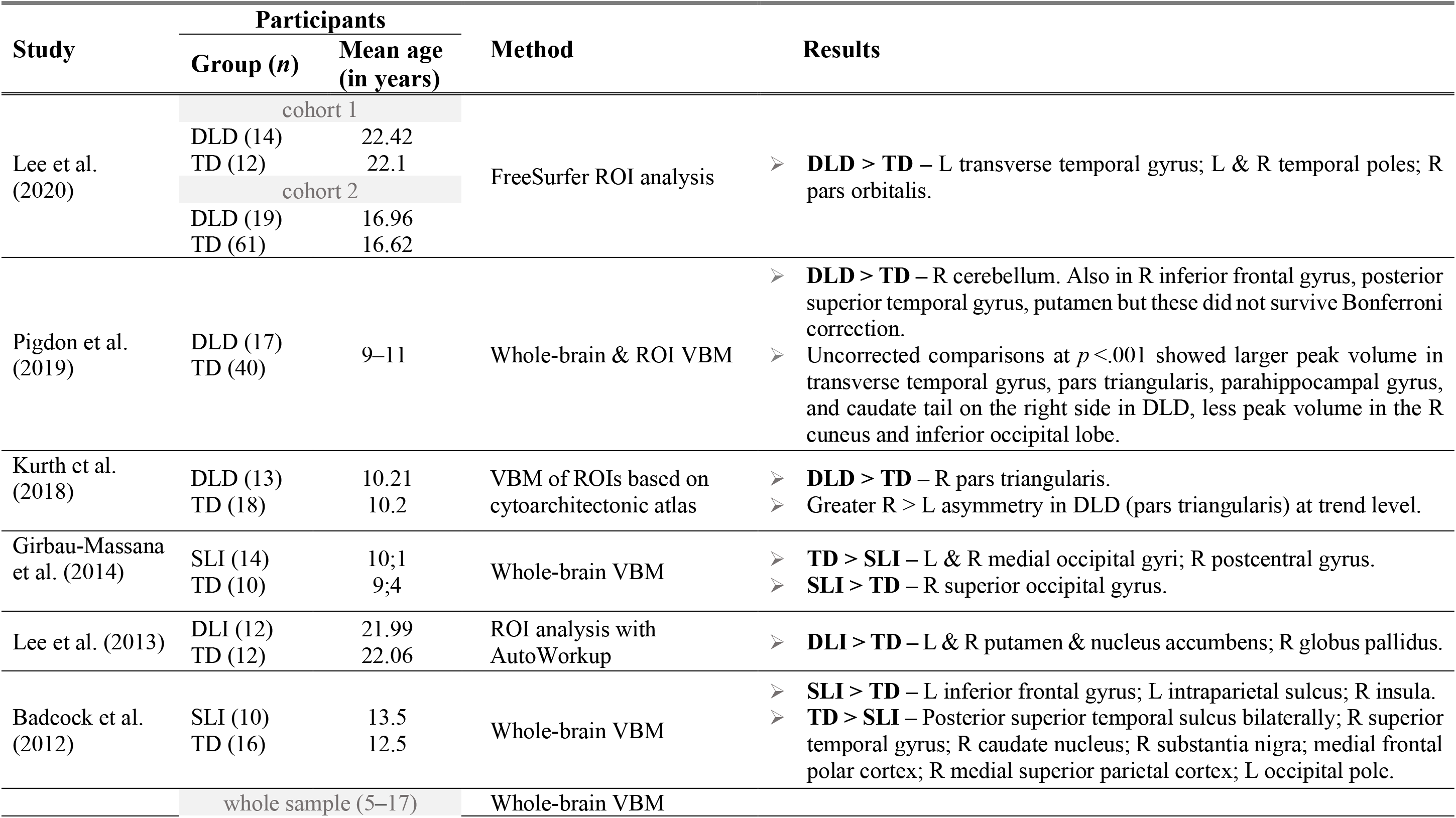
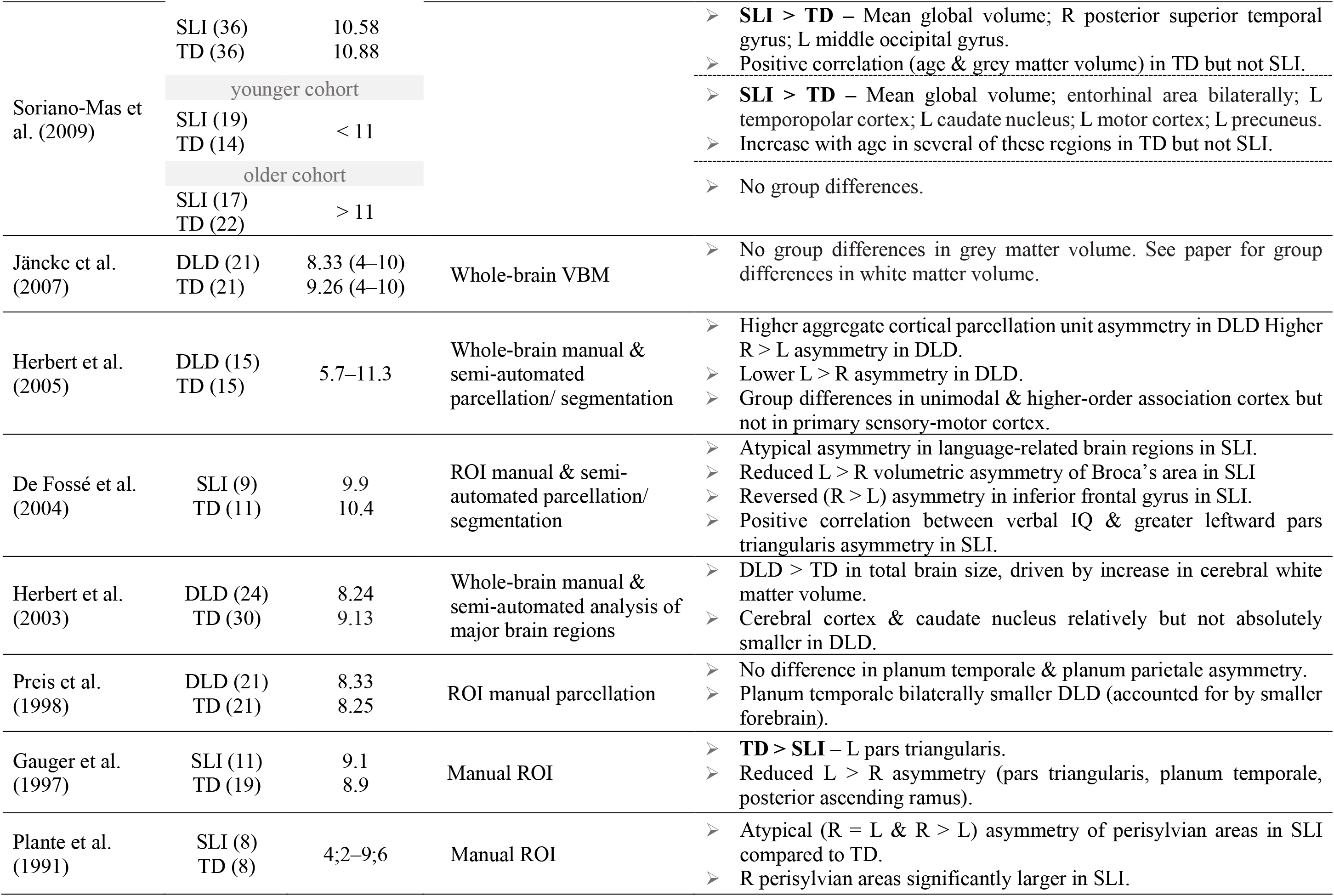
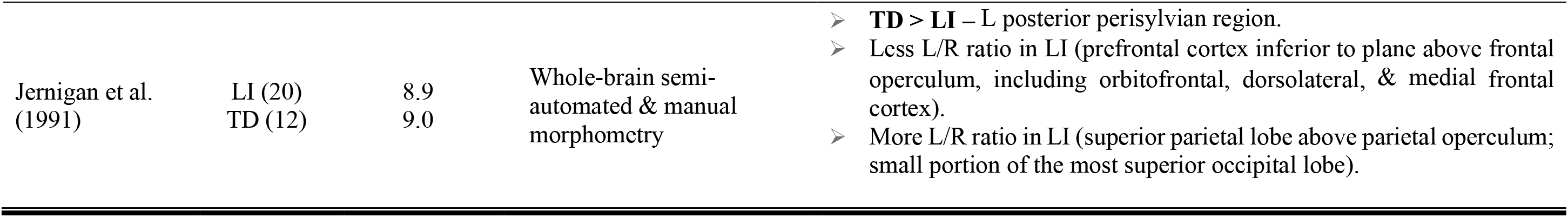
Summary of studies investigating structural grey matter correlates of DLD. The second column provides information of the sample size (only for DLD and controls) and their mean age in years. The age range is provided in parentheses when available. Results pertain to grey matter volume measurements of cortical regions, unless otherwise specified.

In terms of cortical anomalies in the frontal lobes, both lower (Gauger et al., 1997) and greater (Badcock et al., 2012) size of the left inferior frontal gyrus in DLD were reported. Greater volume in DLD was also found in perisylvian regions in the right hemisphere (Plante et al., 1991, Soriano-Mas et al. 2009) including in the pars orbitalis (Lee et al., 2020), pars triangularis (Kurth et al., 2018), and the insula (Badcock et al., 2012). Posteriorly, there are reports of lower volume in DLD in the left (Jernigan et al., 1991) as well as the right hemisphere, comprising the planum temporale (Preis et al., 1998), other portions of the superior and middle temporal gyri, and parietal regions (Badcock et al., 2012; Girbau-Massana et al., 2014). Greater volume in DLD was seen in the transverse temporal gyrus, temporopolar cortex (Lee et al., 2020; Soriano-Mas et al., 2009), medial parietal cortex (Soriano-Mas et al., 2009), and intraparietal sulcus (Badcock et al., 2012). Smaller volume of the right perisylvian region and the occipital petalia were associated with higher verbal intelligence and receptive vocabulary in DLD, respectively (Soriano-Mas et al., 2009).

Atypical structural asymmetry in perisylvian areas was reported in DLD, especially less leftward asymmetry of the inferior frontal gyrus (see Mayes et al., 2015 for a review). More pronounced rightward (i.e. right-greater-than-left) asymmetry of pars triangularis was reported in DLD, and a positive correlation showing that higher verbal intelligence quotient (IQ) scores were associated with more leftward asymmetry of pars triangularis (De Fossé et al., 2004; Kurth et al., 2018). The atypical pars triangularis asymmetry was specifically associated with language difficulties; boys with DLD or with autism spectrum disorder (ASD) and language impairment showed atypical asymmetry, whereas those with ASD but no language impairment did not (De Fossé et al., 2004). Some studies also reported atypical or weaker asymmetry of the planum temporale in DLD (Jernigan et al., 1991; Plante, 1991; Plante et al., 1989, 1991; but see Preis et al., 1998). In a whole brain analysis, children with DLD showed greater rightward asymmetry in the aggregate amount of cortical parcellation units and smaller volume of left-asymmetrical cortex, with significant differences in language-related cortical parcellations, including the posterior supramarginal gyrus and planum polare (Herbert et al., 2005).

An important limitation in past studies is that almost all findings pertain to measures of grey matter volume, which depends upon both surface area and cortical thickness. It remains unclear whether the reported morphological abnormalities reflect differences in surface area, cortical thickness, or both. Surface area and cortical thickness are heritable anatomical features that are under separate genetic influences and have independent developmental trajectories, suggesting that their changes during childhood reflect different developmental processes (Panizzon et al., 2009; Winkler et al., 2010). Investigating surface area and cortical thickness as distinct grey matter volume components may therefore enhance our understanding of the underlying biological processes associated with cognitive functioning in DLD.

Here, we addressed the shortcomings of previous studies by investigating differences in cortical surface area and thickness in a large and well-characterised sample of children with DLD compared with a group of typically developing (TD) controls. FreeSurfer was used to generate mesh models of the cerebral cortex from MRI scans. We also investigated asymmetries in these measures using a reproducible and automated surface-based interhemispheric registration method (see Greve et al., 2013; Sha et al., 2021). Based on the previous literature and behavioural profile, we predicted that relative to TD controls, children with DLD would exhibit: (1) differences in the morphology of perisylvian regions relevant for language, specifically the inferior frontal gyrus; and (2) atypical cortical asymmetry between the same left hemisphere regions and their right hemisphere homologues. We expected morphological differences to include right hemisphere homologues of language regions due to accounts of children with typical language acquisition following focal unilateral brain injury (e.g. Newport et al., 2017), or even without a left hemisphere (Asaridou et al., 2020; Curtiss et al., 2001), indicating the potential of the intact right hemisphere to support language. Accordingly, in DLD where there are no obvious lesions to explain the language impairment, we expect that causal abnormalities in the cortex would occur bilaterally else the high degree of plasticity available during development would support reorganisation of function to the unaffected hemisphere. Furthermore, given DLD’s genetic component (Bishop, 2014), it is likely that genetic disruptions will affect neural development of pathways in both hemispheres rather than only one (e.g. Belton et al., 2003), though the latter possibility cannot be ruled out.

## 2 METHODS

### 2.1 Ethics

Ethical approval was granted by the Medical Sciences Interdivisional Research Ethics Committee at the University of Oxford (R55835/RE002). We obtained written informed consent from the parents/guardians of the participants and written assent from the participants.

### 2.2 Participant eligibility criteria

One-hundred and seventy-five children took part in the Oxford Brain Organisation in Language Development (OxBOLD) project (see Krishnan et al., 2022, 2021). We included participants older than 10 years and younger than 16 who had grown up in the United Kingdom speaking English. Participants were recruited through a range of venues, including those who took part in the SCALES study, (Norbury et al., 2016), the Wellcome Reading and Language Project (Snowling et al., 2015), and the OSCCI Twins study (Wilson & Bishop, 2018). We also recruited from schools for children with language learning difficulties and organisations conducting outreach with those with language impairments (ICAN, Afasic, RADLD), as well as dyslexia (British Dyslexia Association). Typically developing controls were recruited mainly from local schools and schools participating in university outreach programs.

Participants were excluded via parental interview if they (1) had a diagnosis of another developmental disorder (e.g. Down syndrome, ASD, or Attention deficit hyperactivity disorder (ADHD); (2) indicated a history of neurological impairments or neurological disorders such as epilepsy; (3) scored above seven on the Hyperactivity subscale of the Strengths and Difficulties Questionnaire (SDQ; Goodman, 1997); (4) scored above 15 on the Social Communication Questionnaire (SCQ; Rutter et al., 2003); or (5) had any contraindication to MRI. We did not exclude participants based on handedness or speaking multiple languages.

All children passed audiometric screening at 25 dB at 500 Hz, 1000 Hz, and 2000 Hz in the better ear. After testing, we excluded data from 15 children for the following reasons: three obtained a mean non-verbal intelligence quotient (IQ) below 70 (two standard deviations below the mean) based on the composite score obtained from the Block Design and Matrix Reasoning subtests of the Wechsler Intelligence Scale for Children-4^th^ Edition (WISC-IV; Wechsler, 2003); three were later discovered to have not grown up in the United Kingdom speaking English; one participant had scores on two or more tests of language that were more than one standard deviation below the mean but no history of speech and language difficulties; two did not complete the behavioural assessment; three did not have MRI; three were excluded due to incidental findings on the MRI of unknown clinical significance. These exclusions resulted in a cohort of 160 children for whom both MRI and behavioural data were available for analysis.

### 2.3 Behavioural measures

We administered a comprehensive neuropsychological test battery to evaluate the children’s language, memory, non-verbal reasoning, reading, and motor skills. A detailed description of each test can be found in Krishnan et al. (2021). Children were categorized as having DLD if they (1) had a previously reported history of speech-language difficulties, and (2) received a score that was one or more standard deviations below the normative mean on two or more of the six standardized language measures administered. The measures included tests of receptive and expressive vocabulary, sentence comprehension, sentence repetition, and comprehension and recall of narrative. Out of the 160 participants, 55 met the criteria for DLD (*m* = 12.36; 15 females), and 77 were classified as typically developing (TD; *m* = 12.52; 33 females). An additional 28 did not meet the eligibility criteria for DLD at the time of testing but had a history of speech-language problems (HSL; *m* = 12.39; 5 females).

### 2.4 MRI acquisition

Brain scans were acquired on a 3-T Siemens Prisma scanner with a 32-channel radiofrequency head coil. The anatomical images were obtained in the sagittal plane using a T_1_-weighted 3-D MPRAGE sequence with the following pulse sequence parameters: repetition time = 1900 msec, echo time = 3.97 msec, flip angle = 8°, inversion time = 904 msec, with 192 × 192 sagittal field of view, 174 slices, and voxel size = 1×1×1 mm. Participants wore earplugs and noise-cancelling headphones held in place by inflatable air pads. The total image acquisition time was 5 min and 30 sec.

### 2.5 MRI processing

Prior to processing, an experienced rater carefully checked structural scans for artefacts. We used FreeSurfer version 7.2.0 (http://surfer.nmr.mgh.harvard.edu) to automatically parcellate T_1_-weighted images and generate 3-D reconstructions of each participant’s cortex via the “recon-all” command (Dale et al., 1999; Fischl et al., 1999). The reconstructed surfaces were then registered to the *fsaverage* template and assigned a neuroanatomical label (Fischl et al., 1999). To label the sulci and gyri, the Desikan-Killiany atlas was employed, which consists of 33 regions of interest (ROIs) per hemisphere, for a total of 66 ROIs (Desikan et al., 2006; Fischl et al., 2004).

We used the Qoala-T supervised-learning tool (Klapwijk et al., 2019) to assess the quality of cortical segmentations and parcellations post-processing. This automated approach provided a number of advantages over manual correction, including reduced time and variability associated with manual quality control (e.g. inter-rater bias). Four participants were excluded after running Qoala-T (Klapwijk et al., 2019) on the T_1_-weighted images. Qoala-T evaluates scan quality on a 0–100 scale. Nine scans with an average scan quality score below 50 were recommended for exclusion (the mean Qoala−T score of our entire dataset was 71.5). We retained five of these scans after visually inspecting the parcellation qualities. We additionally inspected the quality of image segmentations and cortical reconstructions in 51 scans with average quality above 50, as recommended by Qoala-T. We retained all 51 scans. The final cohort consisted of 54 children with DLD (*m* = 12.43 years; 15 females), 74 TD (*m* = 12.6 years; 33 females), and 28 HSL (*m* = 12.4 years; 5 females).

### 2.6 Cortical metrics

Our morphometric measures of interest were surface area, cortical thickness, and grey matter volume. Surface area is defined as the average area of the triangles (on the tessellated surface) of which each vertex is a member, while cortical thickness is the distance between the white and pial surfaces at each vertex (Fischl & Dale, 2000). Grey matter volume is the product of surface area multiplied by cortical thickness at each vertex.

### 2.7 Whole-brain cortical morphometry

The surface area, cortical thickness, and grey mater volume maps were spatially smoothed in each hemisphere using a Gaussian smoothing kernel with a full width at half maximum (FWHM) of 10 mm. A two-group general linear model (GLM) was performed to investigate cortical differences between DLD and TD in vertex-wise whole-brain analyses. Area, thickness, and volume were separately entered as outcome variables with participants’ age and gender as covariates of no interest. Clusters were defined as groups of contiguous vertices using a vertex-wise threshold of *p* < .001 (two-tailed). We corrected for multiple comparisons by using a cluster-wise correction with a Monte Carlo simulation of 10,000 iterations and a threshold of p < .05 (two-tailed).

Children in the HSL group were excluded from the whole-brain DLD-TD comparisons. Instead, we included them in continuous analyses to examine the effects of age on area, thickness, and volume at each vertex in all participants, with gender as a grouping factor. We then assessed the effect of gender on the cortical metrics (controlling for age) in all participants and the interaction between age and gender.

### 2.8 Surface-based structural asymmetry analysis

Following the steps proposed by Greve and colleagues (2013), we first registered the participants’ surface area, cortical thickness, and grey matter volume maps of both hemispheres to FreeSurfer’s symmetric standard template (*fsaverage_sym*). This step aimed to achieve vertex-wise correspondence between the two hemispheres through precise alignment of the cortical folding patterns between- and within-participants. For each participant, we calculated the asymmetry quotient (AQ) separately for each pair of vertices matched across the left (L) and right (R) hemispheres using the equation:

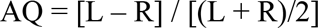

with negative values indicating right- and positive values indicating left-lateralisation. We masked out a region around the boundary between cortical and subcortical structures which had extreme AQ values in many participants, likely due to a warping artifact. We excluded any vertices from all individuals where they had asymmetry values of |AQ| > 1 in at least one quarter (*n* = 31) of our participants (see Supplementary Material, Figure 1). Extreme AQ values were subsequently excluded from the thickness and volume analyses.

**Figure 1.**
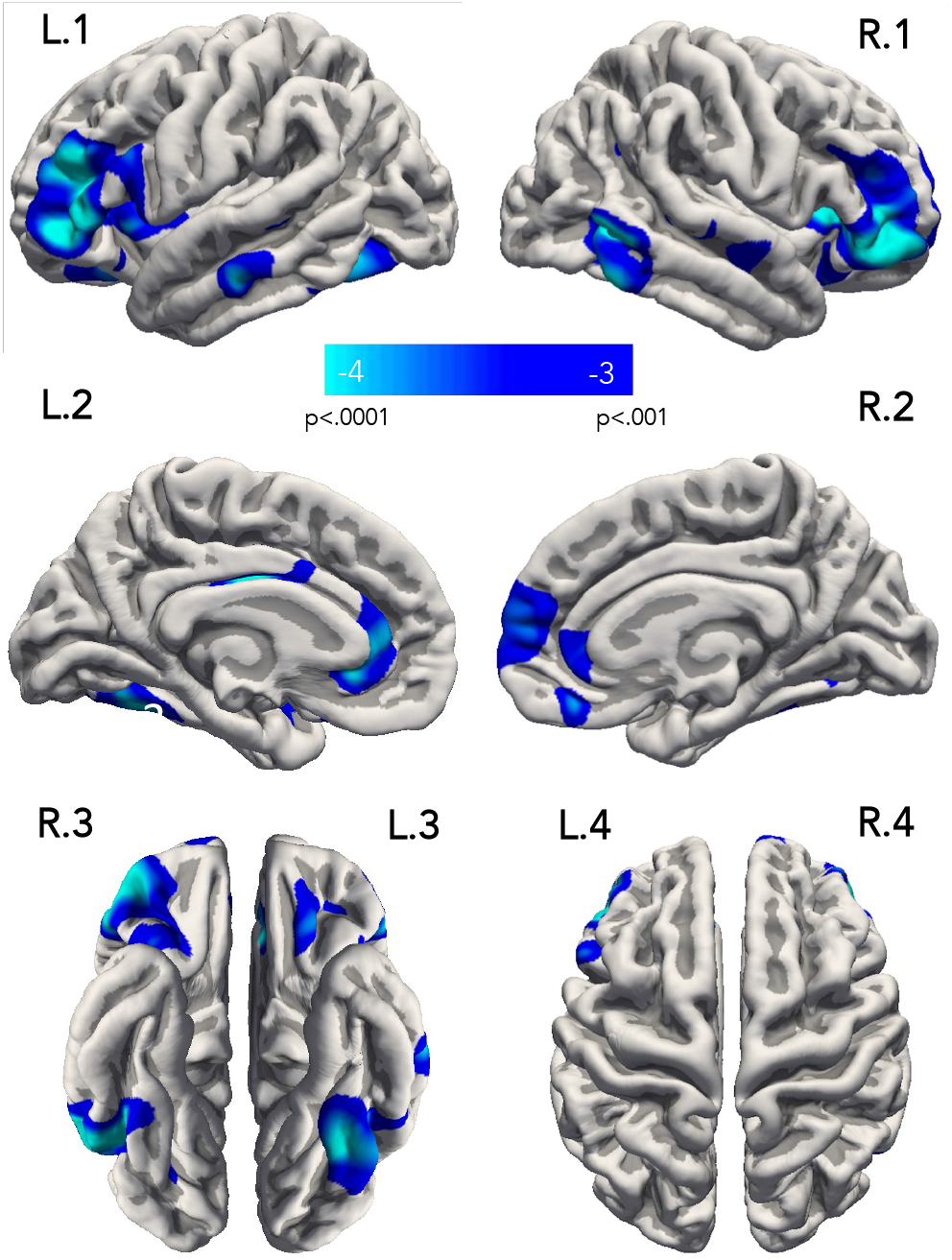
Cortical regions showing smaller surface area in DLD. Whole-brain statistical map illustrating group differences in surface area between DLD and TD children. The left and right hemispheres are shown on the left and right sides of the figure, respectively. The statistical map is overlaid on the pial surface of *fsaverage* and is displayed using an uncorrected vertex-wise threshold of *p* < .001 for visualisation purposes. The numbers inside the colour bar display the −log10(*p*), where *p* is the significance. The blue/cyan hues indicate the DLD < TD contrast. Cluster-corrected results are reported in the text. L.1/R.1 = lateral view; L.2/R.2 = medial view; L.3/R.3 = ventral view; L.4/R.4 = dorsal view.

We used the vertex-wise AQ values between the DLD-TD participants as the dependent variable in a GLM with age and gender as covariates of no interest. Given previous reports of gender differences in grey matter in Broca’s area (Kurth et al., 2018), we also assessed the interaction between group and gender. AQ values from area, thickness, and volume were each entered into the model above. Significant clusters were determined using SurfStat (https://www.math.mcgill.ca/keith/surfstat/) with a cluster-forming threshold of p < .05 (vertex-wise p < .001) and assessed against a random field with the same spatial correlation as the real map to minimise the likelihood of false positives. Clusters identified in this manner had a peak vertex and contiguous vertices, with cluster-wise significance determined.

### 2.9 Brain-behaviour relationships

Comparison of DLD and TD groups on behavioural scores showed that children with DLD had significantly lower scores on tests of non-verbal IQ, expressive and receptive language, reading, memory, and motor function (see Krishnan et al., 2021 for details). Of these we selected non-verbal IQ, non-word repetition, and composite reading scores (from the Test of Word Reading Efficiency (TOWRE); Torgesen, 1999) to test for brain-behaviour relationships in an exploratory analysis. These choices were driven by hypothesised relationships between the brain regions showing differences in the DLD group and did not include any of the six standardised language measures of receptive and expressive vocabulary, sentence comprehension, sentence repetition, and comprehension and recall of narrative that were used to classify the DLD group. We constructed linear regression models to assess whether these behavioural measurements in DLD and TD children could explain the variation seen in morphological values extracted from the statistical clusters showing TD > DLD group differences in surface area in the FreeSurfer analysis. Therefore, we first investigated whether scores on the non-word repetition test correlated with measures from the left inferior frontal regions known to be important for speech production. We also tested whether the word and non-word subtests of TOWRE (Torgesen, 1999) correlated with left inferior temporo-occipital regions implicated in reading development. Non-verbal IQ was included in both sets of analyses to determine whether variance in cortical measures was best predicted by this variable or the variable of interest. To account for multiple comparisons, significance was set at *p* < .016 after Bonferroni correction for three tests.

### 2.10 Data availability

All statistical analyses were conducted using Matlab R2022b (TheMathWorks, Inc) and the R programming language (R Core Team, 2022). The data and code supporting the findings of this study are openly available on the OSF (<u>osf.io/wmkrq</u>). Statistical maps of group differences can be viewed on NeuroVault: https://identifiers.org/neurovault.collection:13977.

## 3 RESULTS

Table 2 presents the demographic information, results from the neuropsychological battery, and gross cortical measures for the DLD and TD groups. There were no significant differences between groups in mean age or distribution of handedness. While the TD group were balanced for gender, there were more males than females in the DLD group, and this group difference was statistically significant (χ^2^ (2, *N* = 128) = 7.88, *p* = .019). The DLD group scored significantly lower than TD on all measures of language, memory, non-verbal reasoning, motor, and reading skills.

**Table 2.**
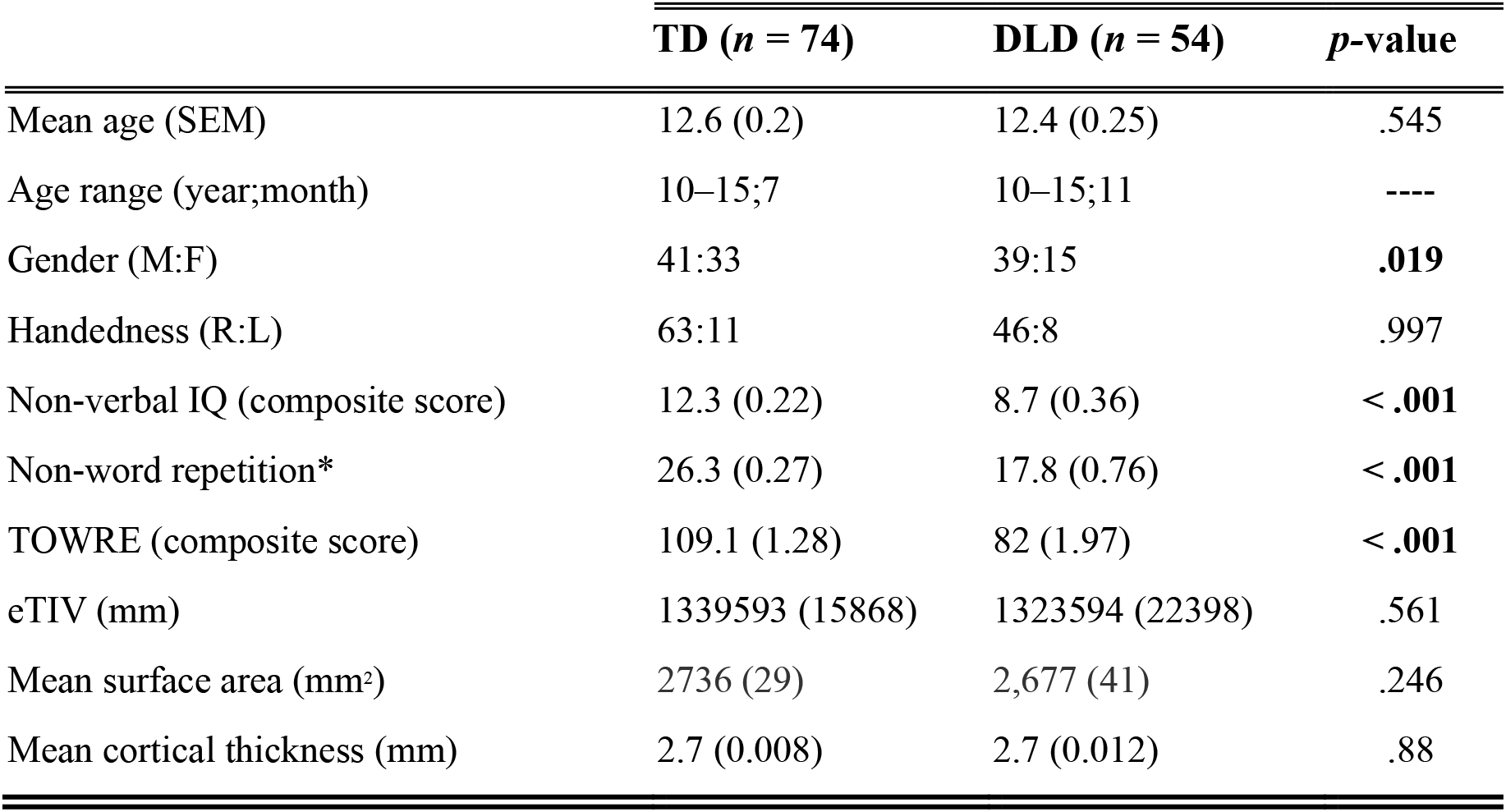
Descriptive statistics for the TD and DLD groups. Descriptive statistics for the TD and DLD groups. The participants’ demographics, including mean age in years, age range (year;month), gender and handedness ratios, estimated total intracranial volume (eTIV), mean whole-brain surface area and cortical thickness estimates, and results of behavioural tasks are presented with standard error of the mean (SEM) in parentheses. Gender and handedness were compared with a chi-squared test. All other variables were compared between groups using a t-test. The last column displays the p-values of these group differences, with those with p < .05 shown in bold typeface. M = male; F = female; R = right; L = left.

### 3.1 Whole-brain analysis of cortical morphology in DLD vs. TD children

The DLD and TD groups did not differ in eTIV (extracted with FreeSurfer’s “aparcstats2table” command), mean surface area and mean cortical thickness (see Table 2). We examined the influence of age and gender on our three cortical metrics (area, thickness, and volume) using the whole cohort (*n* =156; i.e. including the HSL group). For surface area, there was no relationship with age and no effect of gender in any brain region. Regarding cortical thickness, a small number of regions exhibited a decrease in thickness with increasing age, including portions of the medial parietal and occipital cortex, left ventral occipital cortex, and right orbitofrontal cortex. Conversely, increases in thickness with age were detected in the posterior ventro-medial frontal cortex and temporal poles. We found that females had greater thickness than males in the left fusiform cortex only. For volume, we observed a significant negative correlation with age in only a portion of the medial occipital cortex, with no effect of gender. We included age and gender as covariates in the analyses that compared DLD and TD children below (refer to Supplementary Figures 2-3 and Tables 2-3 for summary of significant clusters). Figure 1 depicts vertex-wise surface maps of brain regions where children with DLD had lower surface area than TD children. The DLD group had significantly smaller surface areas in symmetric portions of the anterior inferior frontal gyrus (pars triangularis and pars orbitalis) in both the left and right hemispheres. This difference extended posteriorly to pars opercularis in the left hemisphere and ventrally into the anterior insula bilaterally (Fig. 1, L.1/L.4 and R.1/R.4). Additionally, smaller surface areas were observed in the lateral orbitofrontal cortex on the ventral surface of the left hemisphere (Fig. 1, L.3). On the medial surface (Fig. 1, L.2 and R.2), smaller surface area in DLD was observed in the rostral anterior cingulate cortex bilaterally, anterior medial superior frontal cortex on the right, and posterior cingulate cortex at the boundary with the corpus callosum on the left. Smaller surface area was also seen in DLD in the posterior temporal cortex and ventral occipito-temporal cortex at the level of the mid-fusiform gyrus bilaterally (Fig. 1, L.1/L.3 and R.1/R.3).

**Figure 2.**
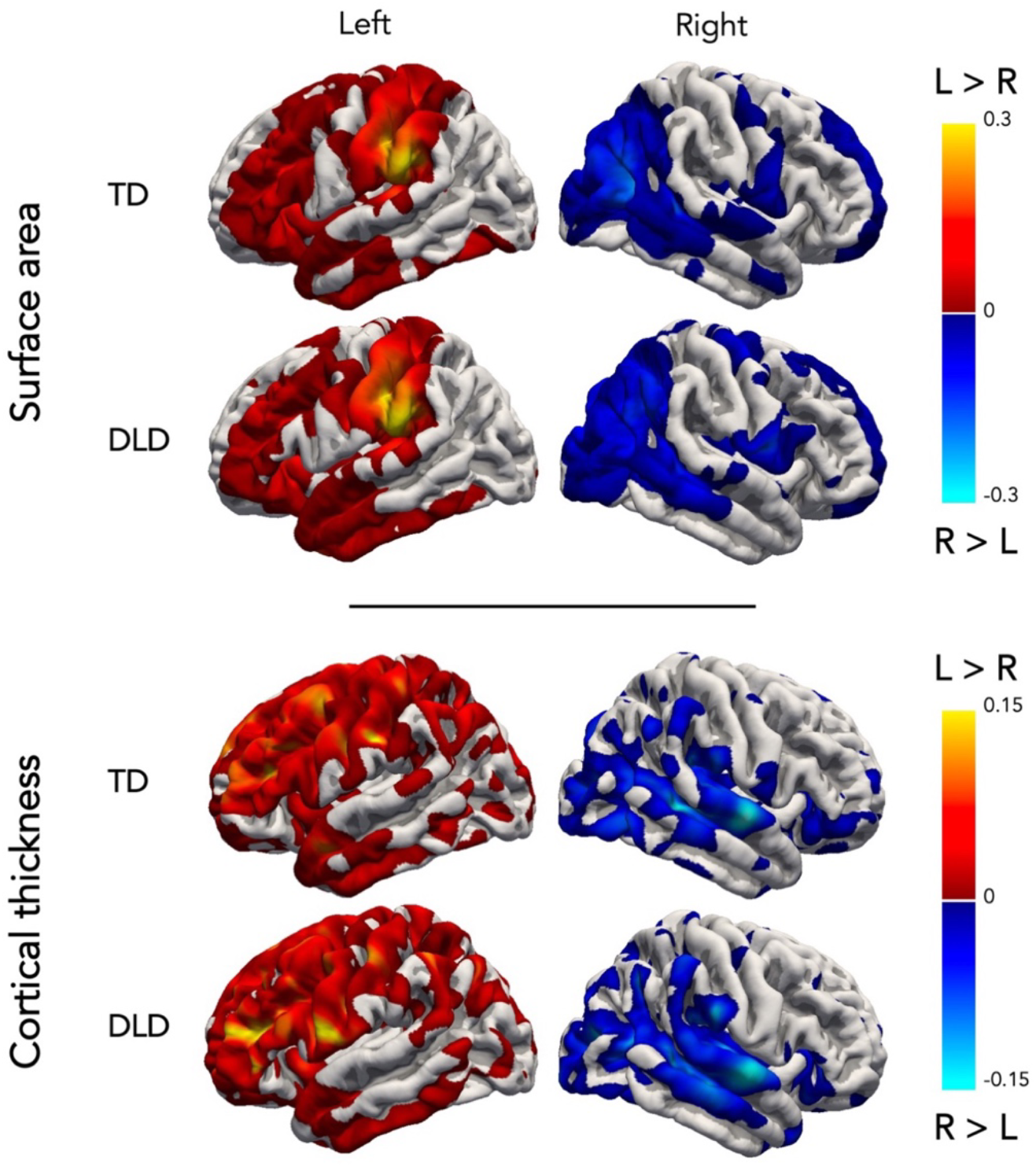
Cortical asymmetries for surface area and thickness in DLD and TD. Whole-brain map of DLD and TD children’s mean asymmetry quotient (AQ) values for surface area and cortical thickness. The map is overlaid on the pial surface (lateral view) and the left and right hemispheres are shown on the left and right sides of the figure, respectively. Red/yellow hues indicate left-greater-than-right asymmetry (L > R), and the blue/cyan hues indicate right-greater-than-left (R > L). L = left; R = right.

**Table 3.** Brain regions showing significant group differences in area and volume. Summary of significant clusters (*p* < .05, corrected) in the DLD < TD comparison for (A) surface area and (B) grey matter volume. The size of each cluster is reported in mm^2^ for surface area and mm^3^ for volume, along with the centroid MNI305 coordinates for Y (coronal), X (sagittal), and Z (axial) planes. The decimal values for the coordinates and cluster size were rounded to the nearest integer.

| Cluster (peak) | Cluster-wise<br>*p*-value | Size | Centroid MNI305<br>Coordinates |  |  |
| --- | --- | --- | --- | --- | --- |
|  |  |  | X | Y | Z |
| (A) Surface area |  |  |  |  |  |
| R superior frontal | .0012 | 617 | 11 | 57 | 10 |
| L pars orbitalis | .0002 | 2269 | -40 | 41 | -6 |
| R lateral orbitofrontal | .0002 | 2212 | 29 | 25 | 3 |
| L rostral anterior cingulate | .0187 | 402 | -61 | 25 | -2 |
| L pars opercularis | .0077 | 471 | -40 | 11 | 9 |
| R posterior temporal | .0002 | 1509 | 56 | -53 | -1 |
| L fusiform | .0002 | 1050 | -33 | -56 | -16 |
| (B) Grey matter volume |  |  |  |  |  |
| R pars triangularis | .0002 | 452 | 47 | 36 | -5 |
| L middle temporal gyrus | .0093 | 245 | -59 | -20 | -22 |
| R posterior temporal gyrus | .0002 | 421 | 49 | -57 | -9 |

Children with DLD also had lower grey matter volume in similar regions to those showing smaller surface area but only small clusters survived thresholding. These were in pars triangularis and the inferior temporal gyrus in the right hemisphere, and the middle temporal gyrus on the left. The anatomical location of statistical peaks using MNI305-space coordinates, along their corresponding *p*-values and the extents of the cluster of voxels (in mm^2^) for surface area and grey matter volume are presented in Table 3. There were no group differences in cortical thickness. The DLD and TD groups did not differ in global surface area or cortical thickness measures (see Table 2).

### 3.2 Analysis of structural asymmetry in DLD vs. TD children

For each of our 54 DLD and 74 TD participants, we generated surface area, cortical thickness, and grey matter volume asymmetry maps in which vertex-wise AQ values across the entire cortex were entered into a GLM with age and gender as nuisance variables. There were no differences between DLD and TD in the degree of structural asymmetry across the cortex in any of these three metrics. Figure 2 shows the mean AQ values indicating the degree of left- and right-ward structural asymmetry in surface area and cortical thickness, superimposed onto the left and right pial surfaces for DLD and TD participants. The patterns in both groups are very similar. In terms of surface area, both groups had a leftwards asymmetry (i.e. left-greater-than-right) in the frontal cortex and a rightwards asymmetry in the occipital lobe. The strongest leftward asymmetries in surface area were located in the supramarginal and post central gyri in both groups. For cortical thickness, there was a leftwards asymmetry in the frontal cortex, which extended posteriorly and dorsally to the parietal cortex. Frontally, the strongest left asymmetries in cortical thickness were seen in rostral middle frontal gyrus, extending posteriorly to pars triangularis and precentral gyrus. The middle to posterior portions of superior and middle temporal gyri showed a rightwards asymmetry in thickness.

### 3.3 Brain-behaviour regression analysis in the whole sample

Non-verbal IQ and non-word repetition scores were entered into linear regression models to identify possible predictors for the mean surface area in the clusters located in pars orbitalis and pars opercularis in the left hemisphere. Non-verbal IQ and the composite TOWRE scores were used in a model to assess whether they predict the mean surface area values in the cluster located in the left ventral occipito-temporal cortex (fusiform).

Visual inspection of the residual QQ-plots showed that the population values were approximately Gaussian for the purposes of building the models. For surface area in the left pars orbitalis, a model with non-word repetition alone (β = .005, *p* = .004) was the best significant predictor of the outcome variable, explaining 5.76% of the variance: *F*(1, 122) = 8.524, *p* = .004. Non-word repetition was also the best single predictor for surface area in the left pars opercularis (β = .004, *p* = .001), explaining 7.18% of the variance: *F*(1, 122) = 10.52, *p* = .001. For surface area in the left ventral occipito-temporal cortex, a model with children’s composite TOWRE scores (β = 001, *p* = .005) was the best significant predictor of the outcome variable, explaining 5.08% of the variance: *F*(1, 122) = 8.142, *p* = .005. Importantly, non-verbal IQ was not a significant predictor of surface area in any of the models.

## 4 DISCUSSION

We investigated cortical morphology in a large, well-characterised sample of 54 children with DLD and 74 age-matched controls aged 10-16 years. Children with DLD had smaller surface area relative to typically developing (TD) controls in the inferior frontal gyrus extending to the anterior insula, in the posterior temporal cortex and the ventral occipito-temporal cortex, all bilaterally. Medially, portions of the anterior cingulate and superior frontal cortex had smaller surface area in DLD. Overall, in terms of anatomical location, these results are consistent with many of the previous findings of cortical differences in DLD reported using other methods and confirm our prediction that children with DLD would show differences in the inferior frontal gyrus. The finding that surface area but not cortical thickness is affected in DLD provides new insights into the underlying processes of brain development in this disorder. To our knowledge, this is the first study in the DLD literature to focus on these separate morphological traits of the cortex. Below we discuss the implications of these findings with relevance to the relationship between cortical structure and function in the context of a developing brain.

### 4.1 Inferior frontal cortex anomalies in DLD

The left inferior frontal gyrus consists of a mosaic of subregions implicated in phonological, semantic, and syntactic processing (e.g., see Vigneau et al., 2006). We found smaller surface area in our DLD cohort in all three subregions of this structure, namely pars orbitalis, triangularis and opercularis, bilaterally. There is only one other study which reported structural abnormality in pars orbitalis in DLD (Lee et al., 2020), but this difference was a volumetric *enlargement* in the right hemisphere and the DLD groups were older on average and smaller than ours (*mean age =* 22.42 and 16.96 years, *n* = 14 and 19 in cohorts 1 and 2, respectively; see Table 1). Two other studies reported volume enlargement in the left inferior frontal gyrus in children with DLD in a roughly similar age range to ours (Badcock et al., 2012; *mean age =* 13.5 years; *n* = 10) and more specifically, in pars triangularis (Kurth et al., 2018, *mean age* = 10.21 years; *n* = 13); but again the sample sizes were much smaller than ours. The only study that reported lower surface area in pars triangularis (measured through manual tracing with a mouse-operated cursor) was by Gauger et al. (1997) in a slightly younger and smaller sample (*mean age* = 9.1 years; *n* = 11).

Pars orbitalis has been associated with an executive function role in semantic cognition tasks in adults (Krieger-Redwood et al., 2015) as well as narrative comprehension (Babajani-Feremi, 2017; Mar, 2010). In children, left pars orbitalis and triangularis are especially implicated in semantic processing (Skeide & Friederici, 2016). It has been argued that children rely more on these regions to process meaning from syntactically complicated sentences while their grammatical skills are still developing (Enge et al., 2021). Pars opercularis becomes a critical hub for syntactic capacity once children’s grammatical skills become more refined towards adulthood (Friederici et al., 2017; Hagoort, 2014). In adults, non-invasive brain stimulation over pars triangularis and pars opercularis selectively impair semantic and phonological judgements respectively (Gough et al., 2005). The strong connections between pars opercularis and the motor representations for the articulators are consistent with its known role in spoken word production (Flinker et al., 2015). Given the functional significance of these cortical regions for language and their substantial connectivity to other brain areas (Bulut, 2022), the anomalous morphology of the inferior frontal cortex in children with DLD is consistent with the behavioural profile of impaired language processing that characterises this population. Accordingly, in an analysis across the TD and DLD children seen in this study, we found that surface area in the left inferior frontal lobe was predicted by performance on a test of non-word repetition.

### 4.2 Ventral occipito-temporal anomalies in DLD

The region showing smaller surface area in the left hemisphere in DLD was very much centred on the mid-fusiform gyrus, the cortical hub for grapho-phonological word processing (Cohen & Dehaene, 2004); furthermore, the size of the surface area was significantly predicted by performance on a test of word and non-word reading. Several studies provided evidence for the involvement of this region in long-term memory representations of visual word forms. Functional and structural abnormalities in this region are reported in children with dyslexia (Kronbichler & Kronbichler, 2018; Paulesu et al., 2014). DLD and dyslexia are distinct neurodevelopmental disorders that frequently co-occur (Bishop & Snowling, 2004) and possibly have a shared aetiology (Ramus et al., 2013). It is therefore not surprising to see overlapping structural abnormalities in both disorders. In fact, prior surface-based morphometric studies have found smaller area in both the inferior frontal gyrus and fusiform gyrus in individuals with dyslexia (Frye et al., 2010).

Good oral language provides the foundation on which later literacy skills develop (Botting et al., 2006). This is supported by studies that found higher risk of reading problems in children with a history of oral language impairments (Snowling et al., 2000). Deficit in spoken language comprehension is a predictor of later reading difficulties in children with DLD (Tallal et al., 1988), likely reflecting the influence of early phonological skills on the development of literacy (Hulme et al., 2015). Cortical differences in the mid-fusiform gyrus in DLD could therefore be attributed to an underdeveloped neural network of skilful reading, perhaps due to lower print exposure.

### 4.3 No differences in cortical asymmetry associated with DLD

Previous structural asymmetry studies linked structural deviation in hemispheric dominance to language impairments (Jernigan et al., 1991; Plante, 1991; Plante et al., 1989, 1991), a finding which we failed to replicate. In fact, we found the pattern of cortical regions showing smaller surface area in DLD to be highly symmetrical. In accordance with a lack of difference in cortical asymmetry, previous work in DLD has also failed to find evidence to support for altered lateralisation of function. For example, a previous functional MRI investigation of children with DLD found no difference in language lateralisation compared with TD during verb generation (Krishnan et al.,, 2021). Additionally, in a well-powered investigation of a sample of twin children with over one-hundred cases of DLD, Wilson and Bishop (2018) used transcranial Doppler sonography (fTCD) to show that there were no differences between school-age DLD and TD children on language lateralisation. In fact, they even found a greater incidence of right laterality for language in TD children, suggesting that an ‘atypical’ pattern of language lateralisation is not necessarily at odds with developing typical language skills (Wilson & Bishop, 2018). On the basis of both our structural findings reported here and previous functional findings, we conclude that there is no evidence for consistently altered language lateralisation or cerebral asymmetry in children with DLD. The inconsistency in current results compared with past findings might either reflect a lack of sufficient statistical power due to the small number of participants, or that previous findings are false positives.

### 4.4 Cortical surface area and thickness development

The differences found in DLD in this study were limited to cortical surface area and grey matter volume, with differences in the latter being neither extensive nor as statistically robust as those seen in the analysis of surface area. There were no group differences in cortical thickness. Since volume is the product of area and thickness, the volume differences were most likely driven by the differences in surface area.

Surface area and cortical thickness are genetically and developmentally distinct components of the cortex (Panizzon et al., 2009; Winkler et al., 2010). Post-natally, surface area increases over the first decade of life and then plateaus during adolescence (Raznahan et al., 2011; Tamnes et al., 2017). In contrast, there are major changes in cortical thickness in the first two years, followed by very slow linear decreases with age that continue across the second decade (Amlien et al., 2014; Lyall et al., 2015). It is worth noting that in our study of children aged 10-16, we found no relationship between age and surface area and only a handful of brain areas showed a reduction in cortical thickness across this age range.

Early in brain development, cortical surface area and thickness are thought to depend on pre- and early post-natal processes of neurogenesis, including the number of radial units, cell proliferation, neuronal migration, programmed cell death, and the density of the columns themselves (Rakic, 2009; Tamnes et al., 2017). In addition, surface area is closely linked to cortical folding, which in turn depends on the division of progenitor cells during early embryonic stages (Chenn & Walsh, 2002; Rakic, 2009). Surface area expansion during childhood is most likely driven by increases in dendritic complexity, spine and synaptic density, and glial cell numbers, along with axonal growth and intra-cortical myelination of these axons. Cortical thickness reductions (or ‘thinning’) may be due to a decrease in synaptic density that arises from experience-dependent plasticity (Rakic, 2009; Sowell et al., 2004), and increased intra-cortical myelin potentially ‘whitening’ the cortex (Paus et al., 2008). Together, these developmental trajectories indicate that cortical volume changes after the first year are most likely due to increases in surface area rather than changes in thickness (Lyall et al., 2015). Cafiero and colleagues (2019) proposed that areal expansion of the cortex during development is related to the myelination of cortico-cortical fibres. Notably, a recent quantitative MRI study provides a clue to differences in the underlying microstructure of the cortex in DLD (Krishnan et al., 2022). Using multi-parametric mapping, these authors found lower levels of myelin, indexed through lower magnetisation transfer saturation values, in DLD in several regions of the left hemisphere’s speech and language network, including in pars opercularis, insula, and several parts of the temporal lobe. In addition, lower mean global values in longitudinal relaxation rate (R1, another myelin marker) were reported for children with DLD relative to TD, reflecting potentially widespread differences in myelin across cortical and subcortical grey matter bilaterally (Krishnan et al., 2022).

Depending on whether cortical differences in DLD are viewed as abnormal or delayed in maturation, lower surface area might be related to any number of the aforementioned neurodevelopmental processes. It might be the case that the affected regions in DLD have not undergone typical processes of neurogenesis pre-natally. Surface area differences between DLD and TD could indeed be driven by genetic risk factors. Animal data has shown that the manipulation of specific genes results in significant changes in the amount of cortical surface area selectively (Mallamaci et al., 2000). Moreover, Beleen and colleagues (2019) found that surface area anomalies in the temporal cortex of a cohort of pre-readers related to a family risk for dyslexia, regardless of their later reading ability.

While surface area differences in DLD might reflect a causal link to genetic disruptions, it cannot be ruled out whether the observed cortical differences are a *consequence* of language learning difficulties. In other words, plastic changes in the cortical areas involved in typical language processing may not have occurred at the same time or rate in the children with DLD, due to their different language-learning experience. The latter argument could explain the strongly left-lateralised pattern of less cortical myelin indexed by magnetisation transfer saturation in DLD in the study by Krishnan and colleagues (2022) compared with the lower R1 myelin marker that affected the brain globally. The language learning impairment seen in our study in children with DLD might have delayed the typical pattern of maturation of cortical areas, through a failure in specialisation of function in circuits involving these regions. Teasing apart the contributions of genetic risk factors and experience-dependent plasticity is an important next step in DLD neuroimaging work, which will require detailed longitudinal investigation.

### 4.5 Limitations & future directions

This study is limited by its cross-sectional nature, which prevents us from discerning whether the observed cortical differences are the cause of DLD or a consequence of having a language disorder. Longitudinal studies of large DLD cohorts in which children are followed over time could shed light on this question, while also having sufficient power to identify statistically robust differences in cortical metrics. Longitudinal investigations could also enhance our understanding of how these structural differences might affect specific aspects of function. For instance, we found a relationship between reading as well as non-word repetition scores with specific brain areas. Pursuing this question is especially important for guiding clinically oriented studies that attempt to answer whether cortical abnormalities could be targeted through behavioural interventions early on, while also gaining insight on the underlying mechanisms of cortical plasticity. Lastly, a longitudinal comparison could also help to understand whether the trajectories of cortical development are abnormal (i.e. deviant from typically developing children at all time points) or delayed (i.e. resemble those of typically developing children at a younger age) in DLD.

Other changes associated with brain development during adolescence complicate attempts to find answers to these questions. Future longitudinal studies with several time points are therefore required to capture the linear and non-linear cortical maturation trajectories in relation to language skills and learn whether the current pattern of finding is unique to the age period studied or is present earlier in development.

## 5 CONCLUSION

Investigating the neural correlates of DLD is particularly challenging given the developmental nature of the disorder and the heterogeneity associated with it. In our investigation of a large and well-characterised DLD neuroimaging dataset, we found that the main cortical difference between typically developing children and those with DLD was in surface area, where a bilateral, and highly symmetric pattern of smaller area was seen in several brain regions in DLD. Notably, these regions survived stringent correction for multiple comparisons. Large DLD cohorts are necessary to identify statistically robust differences in cortical metrics. Future longitudinal work is required to explore how these cortical differences change during development (e.g. whether their trajectory is delayed or abnormal), whether DLD is the cause or consequence of the observed differences, and what their relationship is to function. Clinically oriented studies are also required to assess whether cortical abnormalities could be targeted through training using behavioural interventions early on in development.

## 6 ACKNOWLEDGEMENTS

We are very grateful to our participants and their families for dedicating their time to take part in our study. Without their contribution, this research would not have been possible. We would also like to acknowledge the various individuals and organisations that helped us with recruitment (https://boldstudy.wordpress.com/acknowledgements/). We thank Professor Dorothy Bishop for her encouragement and support. We also thank members of the Wellcome Centre for Integrative Neuroimaging, especially the MRI team at the Oxford Centre for Human Brain Activity: Sebastian Rieger, Juliet Semple, Nicky Aikin, Nicola Filippini, Eniko Zsoldos, and Emily Hinson. We also extend our gratitude to Zhiqiang Sha and Clyde Francks, members of the Language and Genetics Department at Max Planck Institute for Psycholinguistics, for sharing their template scripts for the asymmetry analyses.

