## Supplementary Material for "Differences in cortical surface area in developmental language disorder"

### Contents

Supplementary Figures 1-3: pp. 2-4.

Supplementary Tables 1-3: pp. 6-8.

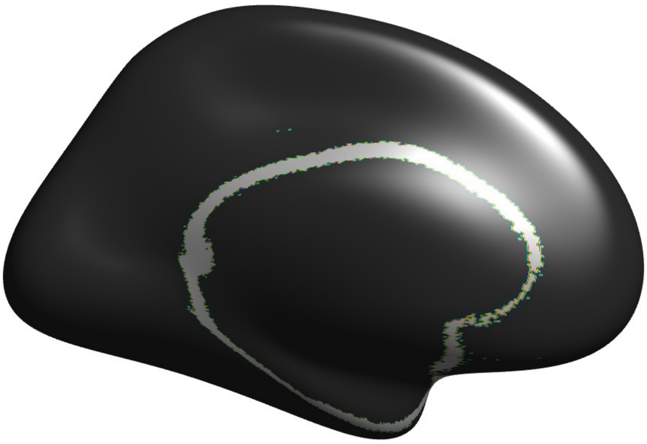

**Supplementary Figure 1.**

The exclusion mask used in the structural asymmetry analysis. The mask comprised vertices at the boundary between cortical and subcortical structures with extreme |AQ| values > 1 in at least one quarter of our participants. This left 157,992 vertex-wise asymmetry measures per individual spanning the rest of the cortex. |AQ| values above one possibly indicate a resampling artifact to the *fsaverage_sym* template.

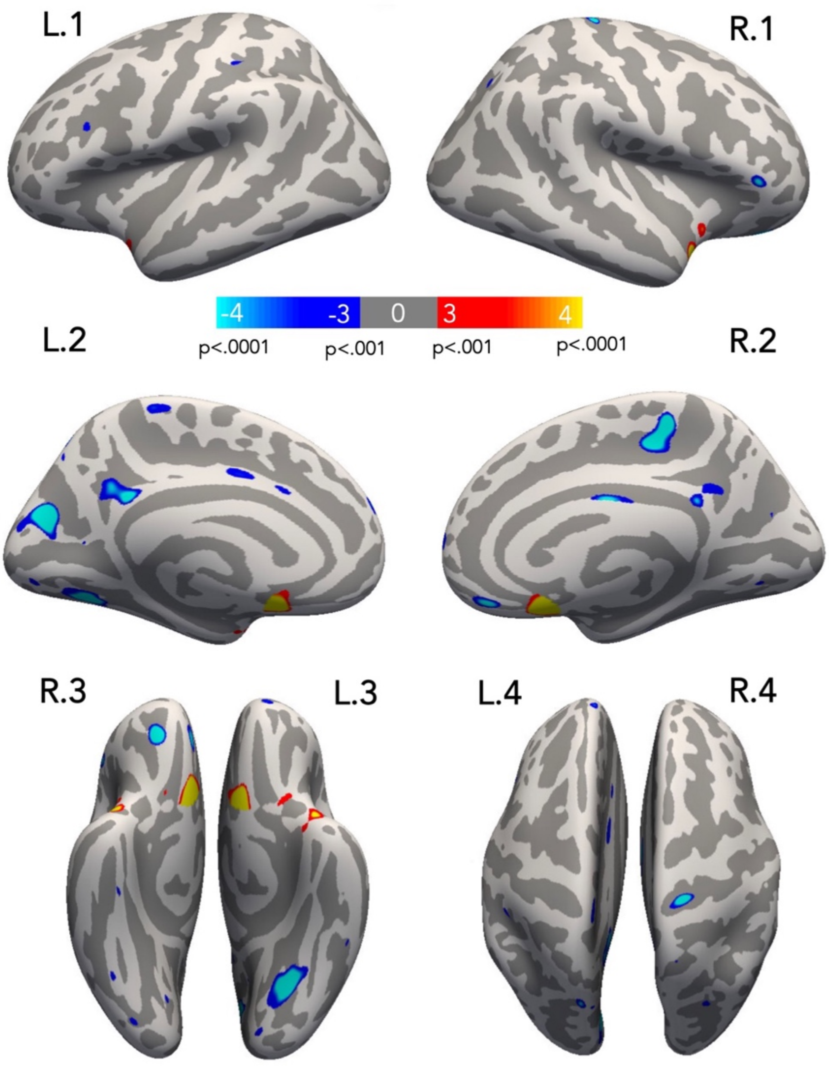


**Supplementary Figure 2.**

Whole-brain significance map of age effects on cortical thickness across all participants, depicted on the inflated surface. Results are displayed using an uncorrected vertex-wise threshold $p < .001$. This is a t-test with, 0 < (DLD+HSL+TD) being positive (red/yellow). The blue/cyan hues indicate regions in which the mean is < 0. Cluster-corrected results are reported Supplementary Table 2. L.1/R.1 = lateral view; L.2/R.2 = medial view; L.3/R.3 = ventral view; L.4/R.4 = dorsal view.

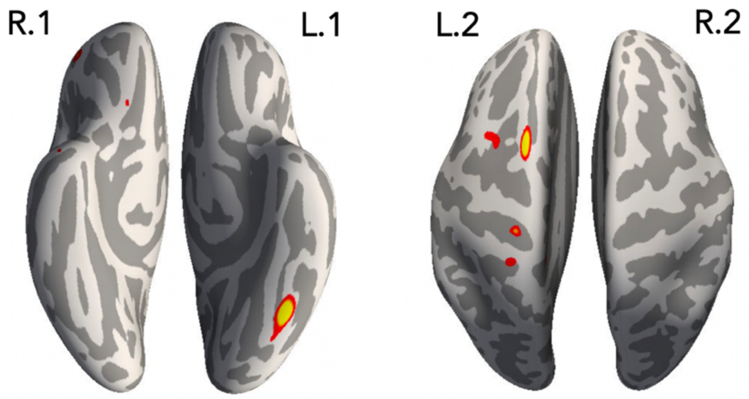


**Supplementary Figure 3.**

Whole-brain significance map of gender effects on cortical thickness across all participants, depicted on the inflated surface. Results are displayed using an uncorrected vertex-wise threshold $p < .001$. This is a t-test with Males < Females being positive (red/yellow). Cluster-corrected results are reported Supplementary Table 3. L.1/R.1 = lateral view; L.2/R.2 = medial view; L.1/R.1 = ventral view; L.2/R.2 = dorsal view.

**Supplementary Table 1.**

Descriptive statistics for the TD, DLD, and HSL groups. The participants' demographics, including mean age in years, age range (year;month), gender and handedness ratios, estimated total intracranial volume (eTIV), mean whole-brain surface area and cortical thickness estimates, and results of behavioural tasks are presented with standard error of the mean (SEM) in parentheses. Gender and handedness were compared with a chi-squared test. All other variables were compared between groups using ANOVA. The last column displays the p-values of these group differences, with those with p < .05 shown in bold typeface. M = male; F = female; R = right; L = left.

| | **TD (*n* = 74)** | **DLD (*n* = 54)** | **HSL (*n* = 28)** | ***p*-value** | **Group differences** |
|---|---|---|---|---|---|
| Mean age (SEM) | 12.6 ± 0.2 | 12.4 ± 0.25 | 12.4 ± 0.29 | .785 | ---- |
| Age range | 10–15;7 | 10–15;11 | 10;5–15;7 | ---- | ---- |
| Gender (M:F) | 41:33 | 39:15 | 23:5 | **.019** | ---- |
| Handedness (R:L) | 63:11 | 46:8 | 24:4 | .997 | ---- |
| Non-verbal IQ | 12.3 ± 0.22 | 8.7 ± 0.36 | 11.3 ± 0.37 | **< .001** | DLD<HSL,TD |
| Non-word repetition* | 26.3 ± 0.27 | 17.8 ± 0.76 | 22.6 ± 0.7 | **< .001** | DLD<HSL<TD |
| TOWRE | 109.1 ± 1.28 | 82 ± 1.97 | 88.1 ± 2.41 | **< .001** | DLD,HSL<TD |
| eTIV | 1339593 ± 15868 | 1323594 ± 22398 | 1409468 ± 34784 | .055 | ---- |
| Mean surface area | 2736 ± 29 | 2677 ± 41 | 2848 ± 62 | **.014** | DLD,TD<HSL |
| Mean cortical thickness | 2.7 ± 0.008 | 2.7 ± 0.012 | 2.7 ± 0.015 | .985 | ---- |

* n = 71 TD, 53 DLD

**Supplementary Table 2.**

Summary of significant clusters for the change in (A) cortical thickness and (B) grey matter volume with age across all participants. The size in $mm^2$ and centroid MNI305 coordinates are reported for significant clusters ($p < .05$, corrected). The coordinate decimals and cluster sizes were rounded to integer values.

| **Cluster (peak)** | **Cluster-wise *p*-value** | **Size** | **Centroid MNI305 Coordinates** | | |
|---|---|---|---|---|---|
| | | | **X** | **Y** | **Z** |
| **(A) Cortical thickness** | | | | | |
| Positive correlations | | | | | |
| L temporal pole | .0431 | 169 | -34 | 5 | -31 |
| R medial orbitofrontal | .0286 | 186 | 6 | 19 | -17 |
| R superior temporal | .0476 | 164 | 41 | 4 | -25 |
| Negative correlations | | | | | |
| L isthmus cingulate | .0201 | 197 | -5 | -45 | 30 |
| L fusiform | .0002 | 452 | -30 | -64 | -6 |
| L cuneus | .0002 | 486 | -12 | -75 | 19 |
| R paracentral | .0084 | 227 | 9 | -33 | 52 |
| **(B) Grey matter volume** | | | | | |
| L cuneus (negative correlation) | .0008 | 362 | -10 | -68 | 10 |

*The extent of each cluster is provided in voxels. Labels of peak location provided by FreeSurfer.*

**Supplementary Table 3.**

Summary of significant clusters in the Males < Females comparison for cortical thickness across all participants. The size in $mm^2$ and centroid MNI305 coordinates are reported for significant clusters ($p < .05$, corrected). The coordinate decimals and cluster sizes were rounded to integer values.

| **Cluster (peak)** | **Cluster-wise *p*-value** | **Size** | **Centroid MNI305 Coordinates** | | |
|---|---|---|---|---|---|
| | | | **X** | **Y** | **Z** |
| **Cortical thickness** | | | | | |
| L fusiform | .0074 | 234 | -42 | -59 | -20 |

*The extent of each cluster is provided in voxels. Labels of peak location provided by FreeSurfer.*
